# AbmR is a mycobacterial dual-function transcription factor and ribonucleoprotein with distinct DNA and RNA-binding determinants

**DOI:** 10.1101/2021.09.03.458936

**Authors:** Roxie C. Girardin, Janice Pata, Xiaohong Qin, Haixin Sui, Kathleen A. McDonough

## Abstract

The bacterium *Mycobacterium tuberculosis* (Mtb) must adapt to myriad host-associated stressors. A recently identified transcription factor, AbmR (ATP-binding *mcr11*-regulator), regulates expression of an essential stress-responsive small RNA (Mcr11) and inhibits the growth of Mtb. Previously, AbmR was found to make 39S complexes of unknown function. Here we report that AbmR 39S complexes are comprised of AbmR and co-purifying RNAs and that RNA-binding inhibits AbmR’s DNA-binding function. While AbmR binds DNA and regulates gene expression in a sequence specific manner, RNA-binding is not sequence specific. Amino acid R146 is important for DNA-binding but completely dispensable for RNA-binding and 39S complex formation, establishing that the RNA- and DNA-binding functions of AbmR are distinct. RNA bound by AbmR was protected from RNase digestion, supporting an RNA modulatory function for the 39S complex. We also found that *abmR* is required for optimal survival during treatment with the ATP-depleting antibiotic bedaquiline, which is associated with extended RNA stability. These data establish a paradigm wherein a transcription factor assembles into large complexes to transition between mutually exclusive DNA-binding gene regulatory and RNA-binding RNA modulatory functions. Our findings indicate that AbmR is a dual-function protein that may have novel RNA regulatory roles in stress adapted Mtb.

## INTRODUCTION

*Mycobacterium tuberculosis* (Mtb) must adapt to a wide array of changing stresses during infection of the human host. Upon infection of macrophages, Mtb establishes a replicative niche within the host cell by subverting normal bactericidal host responses, which allows for the formation of lesions in the host’s lung tissue (1). The vast majority of infections are ultimately arrested, and the bacteria persist within confined granulomas unless reactivation disease occurs. There are many pathways that allow the bacteria to reach a state of non-replicating persistence, which is characterized by metabolic rewiring, low levels of cellular ATP, slow replication, and phenotypic drug tolerance (2-4). Mtb has evolved a complex gene expression regulatory network that includes a wide array of DNA-binding transcription factors, small regulatory RNAs (sRNAs), and stress-responsive signalling cascades that allow for the stress-responsive adaptation Mtb requires to complete its infection cycle (5-9).

A recently identified transcription factor, AbmR (ATP-binding *mcr11*-regulator), regulates expression of an essential stress-responsive small RNA (Mcr11) and inhibits the growth of Mtb (10,11). Expression of *abmR* is up-regulated during nutrient starvation, during macrophage infection, in periods of hypoxia with elevated 3’,5’ cyclic adenosine monophosphate (cAMP), and in response to treatment with the ATP-depleting antibiotic bedaquiline (12-15). The sRNA Mcr11 accumulates in Mtb within the lungs of Mtb-infected mice, contributes to virulence, and has mRNA targets involved in the central metabolism of Mtb (11,16,17). Overexpression of either *abmR* or *mcr11* results in significantly delayed growth kinetics in Mtb (10,18). However, direct targets of AbmR-mediated regulation other than itself and *mcr11* are unknown, as are the DNA-binding sequence determinants of AbmR and the function of the large 39S complexes formed by purified recombinant AbmR.

Here, we report that AbmR 39S complexes are comprised of AbmR and co-purifying RNAs. The RNA-bound state of AbmR was found to compete with AbmR’s DNA-binding function. We identified a tandem, repeating sequence element that is required for AbmR DNA-binding and gene regulatory functions *in vivo*. In contrast, we showed that AbmR’s RNA-binding activity is non-specific and an amino acid residue important for DNA-binding is dispensable for RNA-binding and 39S complex formation. RNA bound by AbmR was protected from RNase digestion and was difficult to displace with a competing RNA ligand. The competition between the DNA-bound and RNA-bound states of AbmR support a physiological function for the 39S complex, which stabilizes associated RNA while decreasing DNA binding. These data establish a paradigm in which a transcription factor can moonlight as a ribonucleoprotein complex that responds to stress in Mtb.

## MATERIAL AND METHODS

### Reagents

Enzymes were purchased from New England Biolabs, USA (DNaseI M0303L, T7 RNA polymerase M0251L, SP6 RNA polymerase M0207S, PacI R0547L, BAMHI R0136L, EcoRI-H R3101L, Murine RNase Inhibitor M0314L, DpnI R0196L, T4 PNK M0201L) or Thermo Scientific, USA (Platinum SuperFi Polymerase 12351050, Phusion polymerase F530XL, DNase and Protease-free RNase A EN0531, TURBO Dnase AM2238)). Nucleic acid purification kits from New England Biolabs were used (Monarch Plasmid Miniprep Kit T1010L, Monarch DNA Gel Extraction Kit T1020S, and Monarch RNA Cleanup Kit T2030L) in combination with phenol:chloroform prepared from molecular grade reagents (Sigma Aldrich, USA), and TRIzol reagent (Thermo Scientific, USA 15596018). Poly(deoxyinosinic-deoxycytidylic) acid sodium salt P4929 and Fluorescein-12-UTP RNA Labelling Mix 11685619910 was obtained from Sigma Aldrich, USA. PCR cloning kits from Thermo Scientific (TOPO TA Cloning Kit 450030) and New England Biolabs, USA (NEB PCR Cloning Kit E1203S) were used for the sub-cloning of DNA fragments and to create plasmid templates for run-off *in vitro* transcription. Ribo Zero rRNA Removal Kit for Bacteria (Illumina, MRZ116C, now discontinued) was used to remove rRNA from *Escherichia coli* total RNA and RNA-seq libraries were constructed using the NEBNext Ultra II Directional RNA Library Prep Kit for Illumina (New England Biolabs, USA E7760L). Libraries were recovered with AMPure XP beads A63880 (Beckman Coulter, USA) and quantified on a Nanodrop (Thermo Scientific, USA) and quality checked with the Agilent High Sensitivity DNA Kit 5067-4626 on a Bioanalyzer (Agilent, USA). Sequencing on the Illumina NextSeq platform was completed using NextSeq™ 500/550 High Output Kit v2.5 (75 Cycles) 20024906 (Illumina, USA). Protein purification was performed using Nickel NTA columns (17-5248-01, GE Healthcare, USA) or beads (7066-4, Millipore,USA). Size exclusion chromatography was completed using a Superdex 200 column (GE Healthcare, USA). RiboReady™ Color 1KB RNA Ladder 89239-528 was obtained from VWR Life Science, USA. Primers used are listed in Supplemental Table 1 and were purchased from Integrated DNA Technologies (USA).

### Biological Resources

Expression strains for the purification of his-tagged AbmR were previously reported (10). Strains used for gene expression analysis in *Mycobacterium tuberculosis* H37Rv (ATCC 25618) were either previously reported or derived from the wild-type sequence parent plasmids previously reported (10).

### Computational Resources

RNA-seq data analysis was completed using Rockhopper freeware using default parameters (19).

### Statistical Analyses

GraphPad Prism version 8 software was used for statistical analyses as indicated.

### Protein Purification and Analysis

Expression and nickel-NTA purification of AbmR was completed as previously described (10). AbmR protein was analyzed by size exclusion chromatography (SEC) on a GE Healthcare Superdex 200 column with a flow rate of 0.5 ml per minute in 50 mM Tris-HCl, 0.25M NaCl, 10% glycerol buffer, pH 8.0. Chromatograms show absorbance at 280 nm and were drawn in Inkscape software. Experiments comparing the impact of DNase I and RNase A on the migration of AbmR by SEC were completed by adding approximately 200 ug of purified AbmR 39S complexes to 20 units of DNase I (New England Biolabs) or 50 ug/ml of protease-free RNase A (Thermo Scientific) to proteins in buffer (50 mM Tris-HCl, 125 mM NaCl, 10% glycerol, 2.5 mM MgCl_2_, 0.5 mM CaCl_2_, pH 8.0) and incubating at 37°C for one hour before analysing the entire reaction mixture by SEC.

For RNase-treated AbmR, approximately 10 mg of purified AbmR 39S complexes were diluted in 10ml of 50 mM Tris-HCl, 10% glycerol buffer, pH 8.0 and bound to 1 ml of Ni-NTA agarose beads (Thermo Scientific) equilibrated in 50 mM Tris-HCl, 10% glycerol buffer, pH 8.0 by gently rocking a sealed disposable gravity column (Thermo Scientific) at 4°C for 1-2 hours. The slurry was then equilibrated at 37°C and 200 ug of DNase and protease-free RNase A and 40 µl of protease inhibitor cocktail (Sigma Aldrich) was added and the column was re-sealed and incubated with gentle rocking at 37°C for one hour. The resin was settled for 10 minutes and unbound protein was allowed to flow through. The bound protein was washed 3 × 10 ml of 50 mM Tris-HCl, 0.75M NaCl, 10% glycerol, 1mM EDTA buffer, pH 8.0. The bound protein was eluted in 9 ml of 50 mM Tris-HCl, 0.75M NaCl, 10% glycerol, 1mM EDTA, 1M imidazole buffer, pH 8.0 and immediately diluted by 50% in 50 mM Tris-HCl, 0.75M NaCl, 10% glycerol, 1mM EDTA, buffer, pH 8.0 to prevent protein precipitation. The protein was dialyzed in a Slide-a-Lyzer cassette with a 3 kDa cutoff (Thermo Scientific) in step-wise gradient of NaCl and imidazole before final dialysis for 16-18h in 50 mM Tris-HCl, 0.25 M NaCl, 10% glycerol buffer, pH 8.0 and concentration by brief dialysis against sucrose. Small aliquots of the protein were flash frozen for further use. The re-purified protein was confirmed to present as a peak with an elution volume like previous RNAse A treated AbmR 39S complexes by SEC and as an intact protein by SDS-PAGE analysis. Quadrupole time-of-flight mass-spectrometry (Agilent) of RNase treated and re-purified AbmR was completed at the University at Albany, SUNY Upstate Proteomics and Mass Spectrometry Core and identified AbmR at the expected molecular mass for an intact monomer. Protein was quantified on a Nanodrop (Thermo Scientific). Negative staining and transmission electron microscopy was completed as previously described with minor modifications (10). Samples were not mixed with gold particles or rinsed and were stained with uranyl formate instead of uranyl acetate (20).

### Nucleic Acid Extraction and Native Gel Electrophoresis

0.5-1 mg of purified AbmR 39S complexes were mixed 1:1 with 25:24 phenol (pH8.0):chloroform and the aqueous phase was precipitated with 3 M sodium acetate, pH 5.3 and 100% ethanol. The precipitated nucleic acid was washed twice with 75% ethanol and resuspended in nuclease free water before being quantified on a Nanodrop (Thermo Scientific). Approximately 2 µg of AbmR-extracted nucleic acid, dsDNA PCR product, or yeast tRNA (Sigma Aldrich) was set up in a reaction series that included mock (buffer), 3 units of DNase I, or 15 µg of RNase A and incubated at 37°C before visualization on a 1.7% agarose gel made up in 0.5X TBE, pH 9.0 and run at 110v for 1.1 hours. Gels were post-stained with ethidium bromide and visualized under UV transillumination on a GelDoc (BioRad Laboratories). Approximately 6 µg of purified AbmR complexes was similarly treated with mock, DNase I, or RNase A and resolved in the same gel. Results are representative of three independent experiments.

### Electromobility Shift Assays with a DNA Ligand

Electromobility shift assays (EMSAs) with AbmR 39S complexes or with de-complexed protein were completed as previously described with some modifications (10). DNA oligos without FAM labels were end-labelled with ^33^P end-labelled DNA probes (0.05 pmol) were incubated with 1.5 µM recombinant AbmR with or without 500-fold excess cold competitor DNA fragments. DNA oligos with a 5’6-FAM label were annealed and used at a concentration of 10 nM in subsequent DNA binding reactions. DNA binding reactions were comprised of 10 mM Tris-HCl pH 9.0, 50 mM KCl, 1 mM EDTA, 50 µg ml–1 BSA, 1 mM DTT, 0.05% NP-40, 20 µg/ml–1 poly dI-dC, 10% (vol/vol) glycerol, 10 nM dsDNA ligand, and 1 mM ATP unless otherwise noted. AbmR protein was added at varying concentrations and the reaction mixture was incubated at room temperature for 30 minutes. When comparing Wt AbmR to AbmR mutant proteins, reaction mixtures contained 0.45 µg to 1.35 µg protein per reaction. DNA-binding reactions were resolved on a non-denaturing 8% (weight/vol) polyacrylamide gel run for 2.5 h, 14 V/cm at 4°C in 0.5X Tris-borate-EDTA (TBE), pH 9.0 running buffer and visualized using the FITC channel set at 100 µM resolution in low intensity mode on a GelDoc Pharos imager (BioRad Laboratories). ImageQuant software was used to analyse the fraction of probe bound in each lane and GraphPad Prism version 8 software was used to calculate the Kd, and Hill slope for AbmR dimers and complexes using a one-site specific binding with Hill slope non-linear regression model.

### Site-directed Mutagenesis

A modified version of Strategene’s QuickChange protocol was used to introduce mutations into the AbmR binding sequence in transcriptional reporter constructs for use in *Mycobacterium tuberculosis*. Parent plasmids of promtor:GFPv fusion constructs pMBC2069 (mcr11 promoter) and pMBC1903 (abmR promoter) were diluted and 15 ng was added to a PCR reaction containing 125 ng of each primer, either Phusion polymerase or SuperFi polymerase (Thermo Scientific) and GC enhancer, and thermocycled (95°C 30s, 61°C 1m, 68°C 7m) for 16 cycles. The enzyme DpnI was added to PCR reactions and parent plasmid was digested at 37°C for four hours before heat-killing the enzyme for 10 minutes at 65°C. Reaction mixtures were directly transformed into chemically competent DH5α *Escherichia coli* and transformants were selected on kanamycin containing LB agar plates. Introduction of the desired mutation was confirmed by Sanger sequencing before transformation of mutant constructs into Rosetta™ (DE3)pLysS E. coli (Novagen) for expression of recombinant protein or *ΔabmR* Mtb for transcriptional activity assays.

### Transcriptional Reporter Assays in Virulent Mtb

*AbmR* or *mcr11* promoter:GFPv reporter fusion plasmids were sequence verified and used to transform *ΔabmR* Mtb (Girardin et al, 2018) by electroporation. Positive transformants were selected by their kanamycin drug resistant phenotype and confirmed by PCR. Low passage stocks were frozen for further experimentation. Middlebrook 7H9 liquid medium (Difco) supplemented with 10% (vol/vol) OADC, 0.2% (vol/vol) glycerol, 10% (vol/vol) and 0.05% (vol/vol) Tween-80 (Sigma-Aldrich) was inoculated with equal amounts of an empty vector negative control, a positive control, or constructs bearing Wt or mutant copies of the reporter fusion constructs and incubated in shaking, hypoxic conditions (1.3% O2, 5% CO2).

At mid-log and stationary phases of growth, aliquots of Mtb cultures were collected and sonicated briefly using a Virsonic 475 Ultrasonic Cell Disrupter with a cup horn attachment (VirTis Company) and chilled, circulating water before duplicate samples were diluted in fresh media in a 96 well plate. The OD at 620 nm was read using a Tecan Sunrise® microplate reader. The level of fluorescence from the GFPv reporter gene was detected using the CytoFluor Multi-Well Plate Reader Series 4000 (PerSeptive Biosystems) at 485 nm excitation and 530 nm emission. Fluorescence levels were normalized to 10^6^ bacteria as determined by OD at 620 nm. Fluorescence of the various mutant reporter constructs were compared Wt levels and the statistical significance of differences between strains was calculated using one-way ANOVA with Sidak’s correction for multiple comparisons in GraphPad Prism version 8 software. Exact p-values are reported on the graph. Ratios are the mean technical duplicates from three independent experiments.

### RNA-seq of AbmR Co-Purifying RNA

*E. coli* bearing the wild type AbmR expression plasmid was induced for AbmR expression, and AbmR was purified on Ni-NTA resin and then phenol:chloroform extracted and ethanol precepted as described above. A duplicate culture of induced *E. coli* was lysed, and total RNA was purified in TRIzol reagent. Approximately 1 µg of purified RNA was treated with TURBO DNase (Thermo Scientific, USA) per the manufacturer’s protocol, and the RNA was isopropanol precipitated after enzyme inactivation. Total RNA from the induced *E. coli* control sample was rRNA depleted using the RiboZero for bacteria kit (Illumina, USA). 100 ng of two replicates each of AbmR co-purified RNA and rRNA-depleted induced *E*.*coli* control RNA were converted into RNA-seq libraries using the NEBNext Ultra II Directional RNA Library Prep Kit for Illumina (New England Biolabs, USA) per the manufacturer’s instructions. Libraries were submitted to the Advanced Genomic Technologies Core at Wadsworth Center for quantitation, quality control, and sequencing on the Illumina Next-Seq platform. Resulting reads were mapped to the *E. coli* BL21(DE3)pLysS Gold genome and the AbmR expression plasmid sequences using Rockhopper freeware with default parameters. All replicates resulted in at least 5 million mapped reads per sample, and RNA-seq traces demonstrated a clear separation of plus and negative strand transcripts. Statistical analysis was performed by Rockhopper. A subset of AbmR-associated RNA transcript sequences with greater than an expression level of 20, significant q-value (<0.005), and >5-fold enrichment in AbmR pull-downs over input controls, or transcripts that represented the top 10-25% of the most abundant reads in the AbmR-associated samples were selected for further follow-up and biochemical characterization.

### Electromobility Shift Assays with an RNA Ligand

RNA that co-purified with Wt AbmR 39S complexes was extracted and purified as described above. Approximately 2 µg of de-complexed AbmR were tested for binding of the purified RNA with or without 1 mM ATP by EMSA. For RNase protection experiments, extracted, co-purifying RNA was added to increasing amounts of RNase A (5-20 µg) in the presence or absence of approximately 2 µg of de-complexed AbmR in a 20 ul reaction. Protein-only controls without RNA ligand were also loaded. Reaction mixtures contained approximately 1.5 µg of AbmR-extracted nucleic acid, 20 units of murine RNase inhibitor (New England Biolabs), 12.5 mM DTT, 10 mM Tris-HCl pH 9.0, 50 mM KCl, 10 mM MgCl_2_, 125 mM NaCl, and 7% glycerol. Reactions were incubated at 37°C for 10 minutes (RNA binding with ATP experiments) or for 5 minutes before the addition of varying amounts of RNase A (0-0.02 µg per 20 µl reaction) and an additional 5 minutes of incubation 37°C before visualization on a 1.7% agarose gel made up in 0.5X TBE, pH 9.0 and run at 110 V for 1.1 hours. Gels were post-stained with ethidium bromide and visualized under UV transillumination on a GelDoc (BioRad Laboratories) and results are representative of minimum of three independent repeats.

The open reading frames of several genes found to have AbmR co-purifying transcripts were sub-cloned into the pMiniT 2.0 and Sanger sequenced to verify insert integrity and orientation. Purified plasmids were linearized with PacI or BamHI and re-purified for use as templates for in vitro transcription with T7 or SP6 RNA polymerases. *In vitro* transcribed RNAs were purified with an RNA Cleanup Kit (New England Biolabs, USA) and quantitated on a Nanodrop (Thermo Scientific). Approximate size as confirmed in a 1.7% agarose gel made up in 0.5X TBE, pH 9.0 and run at 110v for 1.1 hours with a denatured, single-stranded RiboReady 1 kB RNA ladder (VWR Life Sciences, USA). Internally labelled RNA was similarly created using *in vitro* transcription, with Fluorescein-12-UTP labelling mix (Sigma Aldrich, USA). Approximately 1.5 µg of unlabelled RNAs were added to binding reactions for visualization on a 1.7% agarose gel as described above. Labelled RNAs were added to binding reactions with 0.6 (-/+ ATP experiments) - 1 µg (competition and protection experiments) de-complexed AbmR in binding buffer (10 mM Tris-HCl pH 9.0, 50 mM KCl, 10 mM MgCl_2_, 5% glycerol, 200 mM NaCl, 0.2 µg poly-dI-dC, 1 nM yeast tRNA) at a final concentration of 05.-1 nM and incubated at 37°C for 10 minutes before visualized on a 5% 80:1 acrylamide gel made in 0.5X TBE, pH 9.0 and run at 150 V for 4 hours. For competition reactions, the indicated excess of unlabelled AbmR binding site dsDNA or *in vitro* transcribed orthologous RNA was added to the reaction master mix, and then protein was added. For RNase protection experiments, yeast tRNA and poly dIdC was substituted for 20 units of murine RNase inhibitor (New England Biolabs), 12.5 mM DTT, NaCl was decreased to 62.5 mM, and varying amounts of RNase A (0-0.02 µg per 20 µl reaction) were added.

### Bedaquiline treatment of Mtb

*AbmR* or *mcr11* promoter:GFPv reporter fusion strains were grown until late log phase shaking, in ambient air and the OD at 620 nm was read using a Tecan Sunrise® microplate reader. Cultures were diluted in fresh media to an OD_620_ of 0.5/ml and then treated with 25 µg/ml bedaquiline, 7.3 µM isoniazid, or DMSO vehicle control and incubated at 37°C (15). At 4h and 24h post-treatment, aliquots of Mtb cultures were collected and promoter:GFPv reporter fusion expression was measured as described above.

Wt and *ΔabmR* or *ΔabmR* complemented strains of Mtb were grown until late log phase shaking, in ambient air and the OD at 620 nm was read using a Tecan Sunrise^®^ microplate reader. Cultures were diluted in fresh media to approximately 1 × 10^7^ CFU/ml and then treated with 25 µg/ml bedaquiline or DMSO control and incubated at 37°C in ambient, shaking conditions. At the indicated time points, aliquots of each strain/treatment combination were taken, sonicated, and plated for survival on Middlebrook 7H10 + cycolohexmide + OADC agar plates. CFUs were enumerated after 4 weeks of incubation at 37°C. Relative survival was calculated by dividing the CFUs at each time point by input CFUs at t=0 and ratioing to Wt bedaquline treated samples. The results are the means of three independent repeats. A two-way ANOVA with Bonferroni correction for multiple comparisons was used to assess significance.

## RESULTS

### AbmR 39S complex formation is mediated by RNA-binding

While many bacterial transcription factors bind DNA as dimers or tetramers and form condensed nucleoid-associated protein complexes of protein and DNA, the large soluble 39S complex formed by AbmR is unusual (10). We speculated, due to the high A260/A280 ratio of purified AbmR 39S complexes, that the AbmR protein was co-purifying with nucleic acid. Purified AbmR 39S complexes were extracted in phenol:chloroform and co-purifying nucleic acids were ethanol-precipitated from the aqueous phase. The extracted nucleic acid was aliquoted and tested for degradation by DNase I or RNase A and analysed alongside relevant controls (Figure 1A). The nucleic acid that co-purified with AbmR was heterogenous in size and susceptible to RNase A degradation but not DNase I degradation (Figure 1A). Furthermore, purified AbmR 39S complexes were stained by ethidium bromide, and UV signal from the stain was lost when purified AbmR 39S complexes were treated with RNase A but not DNase I. The results demonstrate that AbmR co-purifies with RNA when over-expressed in *E. coli*.

**Figure 1:**
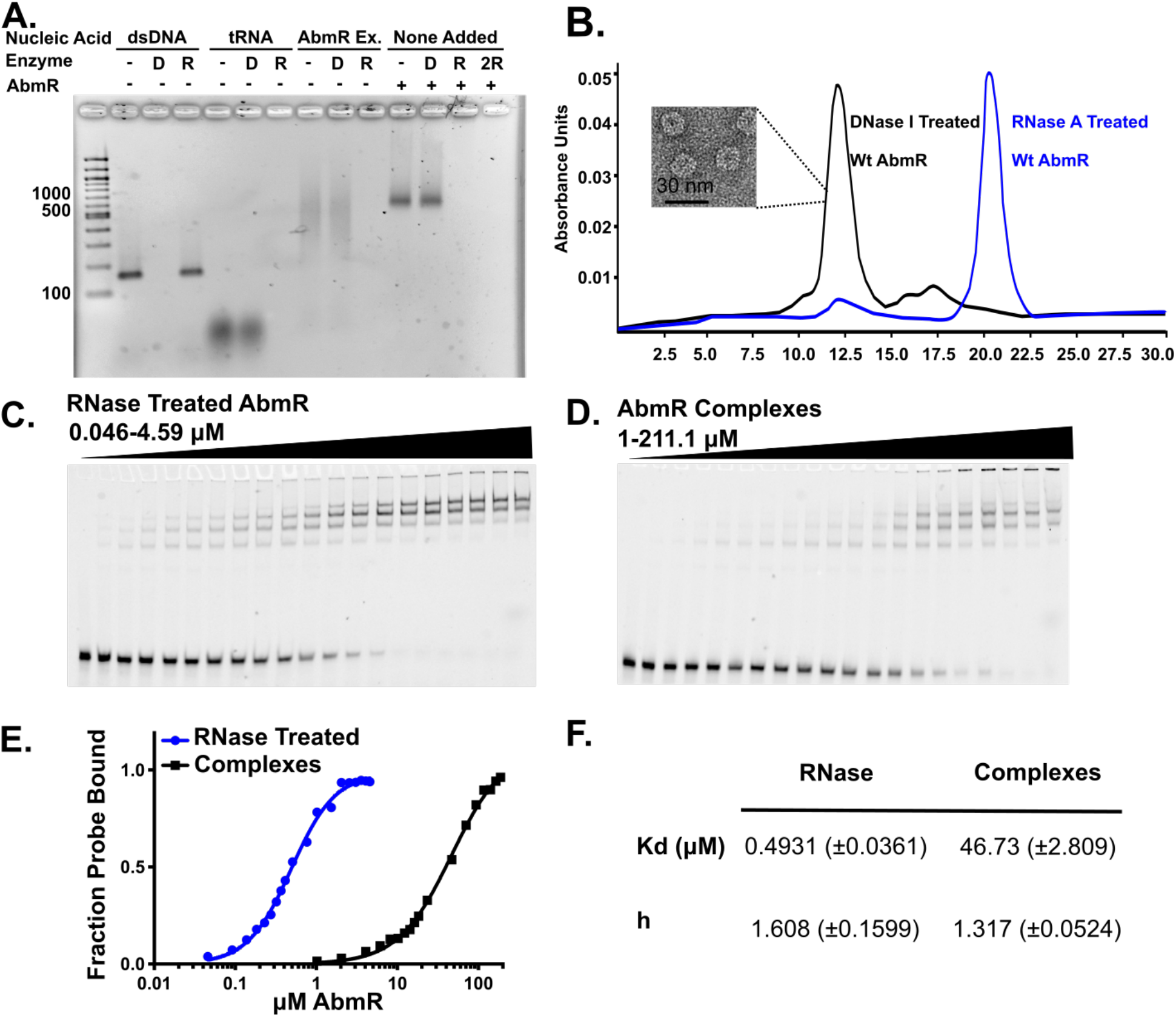
AbmR 39S complex formation requires RNA binding. Formation AbmR ribonucleoprotein complexes is mediated by RNA-binding, in competition with the DNA-binding activity of AbmR. A. Ethidium bromide stained agarose gel of the DNase (D) or RNase (R) digestion of double-stranded DNA (dsDNA), tRNA, phenol-chloroform nucleic acid extracted from purified AbmR 39S complexes, (AbmR Ex.) or AbmR 39S complexes with no added nucleic acid (None Added). A 100 base pair dsDNA ladder is loaded for comparison. C. Size exclusion chromatography of RNaseA or DNAse treated AbmR complexes. Inset shows an electron micrograph of negatively stained AbmR 39S complexes. C. De-complexed AbmR tested for binding of the *mcr11/abmR* promoter DNA fragment with ATP by electromobility shift assay (EMSA). D. AbmR 39S complexes tested for binding of the *mcr11/abmR* promoter DNA fragment with ATP by EMSA. E. Binding plot of the experimental results in (C) and (D). F. Details of the Kd and Hill slope for RNase treated AbmR and 39S complexes using a one-site specific binding with Hill slope non-linear regression model in Graphpad Prism version 8 software.

Next, we evaluated the impact of extensive RNase A digestion on the oligomeric status of purified AbmR 39S complexes and compared it to extensive DNase I digestion using size exclusion chromatography (SEC). RNase A digestion resulted in the near-complete disappearance of 39S complexes and the appearance of a lower molecular weight, de-complexed species (Figure 1B). DNase I digestion had no significant impact on the abundance of AbmR 39S complexes (Figure 1B). We re-purified RNase A-treated, de-complexed AbmR protein and found it presented as an intact mass of the expected size (Supplemental Figure 1A), confirming that the dissociation of the 39S complex was not explained by proteolytic cleavage. Thus, the formation of AbmR 39S complexes is mediated by the RNA-binding activity of the protein.

### RNA-binding inhibits AbmR’s DNA-binding activity

We speculated that the RNA-bound state of AbmR would impact the DNA-binding activity of AbmR if the protein had a single nucleic acid binding domain, or if the RNA-bound state allosterically regulated/physically occluded the DNA-binding activity of AbmR. Electromobility shift assays (EMSAs) were performed with a dsDNA probe containing the sequence upstream of and between the *mcr11* and *abmR* genes. Results of these assays were used to calculate the apparent dissociation constants (Kd) of purified AbmR 39S complexes and de-complexed (RNase A-treated and re-purified, lower molecular weight species) AbmR in the presence of ATP ((10) and Figure 1C-D). De-complexed AbmR was found to have an apparent Kd nearly two logs lower than that of 39S complexes (0.49 uM versus 47 uM) (Figure 1E-F). Both forms of the protein give Hill coefficients greater than 1, indicating that AbmR binds DNA with positive cooperativity. The large difference in apparent Kds of RNA-bound complexes and the de-complexed AbmR shows that RNA-binding inhibits AbmR binding to DNA.

### DNA-binding by AbmR is sequence specific and required for gene regulation

Cooperative binding of DNA by transcription factors occurs when binding to one site facilitates binding at a neighboring site. AbmR:DNA gel shifts present as multiple bands ((10), Figure 2A), suggesting that the *mcr11-abmR* locus binding region contains multiple smaller binding sites. We narrowed the region of AbmR’s binding sequences using scanning competition EMSAs, where excess unlabelled dsDNA fragments with sequence replacements in a sliding window could compete for binding of the labelled fragment (Supplemental Figure 1B-E). The *mcr11-abmR* locus has regions of unusually high (>75%) AT content for mycobacteria (typically <35% AT), so we used both GC-rich and AT-rich replacement sequences to test the specificity of the replacements (Supplemental Figure 1B-D). AbmR’s binding region was localized to Fragment C, which contains two repeating sequence elements: four direct tandem repeats of ATGN_5_ and two spaced-out repeats of ATTA(A) which can be direct or inverted (Supplemental Figure 1E, Figure 2A). Replacement of the ATTA(A) repeats had a modest effect on AbmR’s ability to bind DNA, but resulted in an altered band shift pattern. However, replacement of the four direct tandem repeats of ATGN_5_ (4x ATGN_5_) abolished DNA binding (Figure 2A).

**Figure 2:**
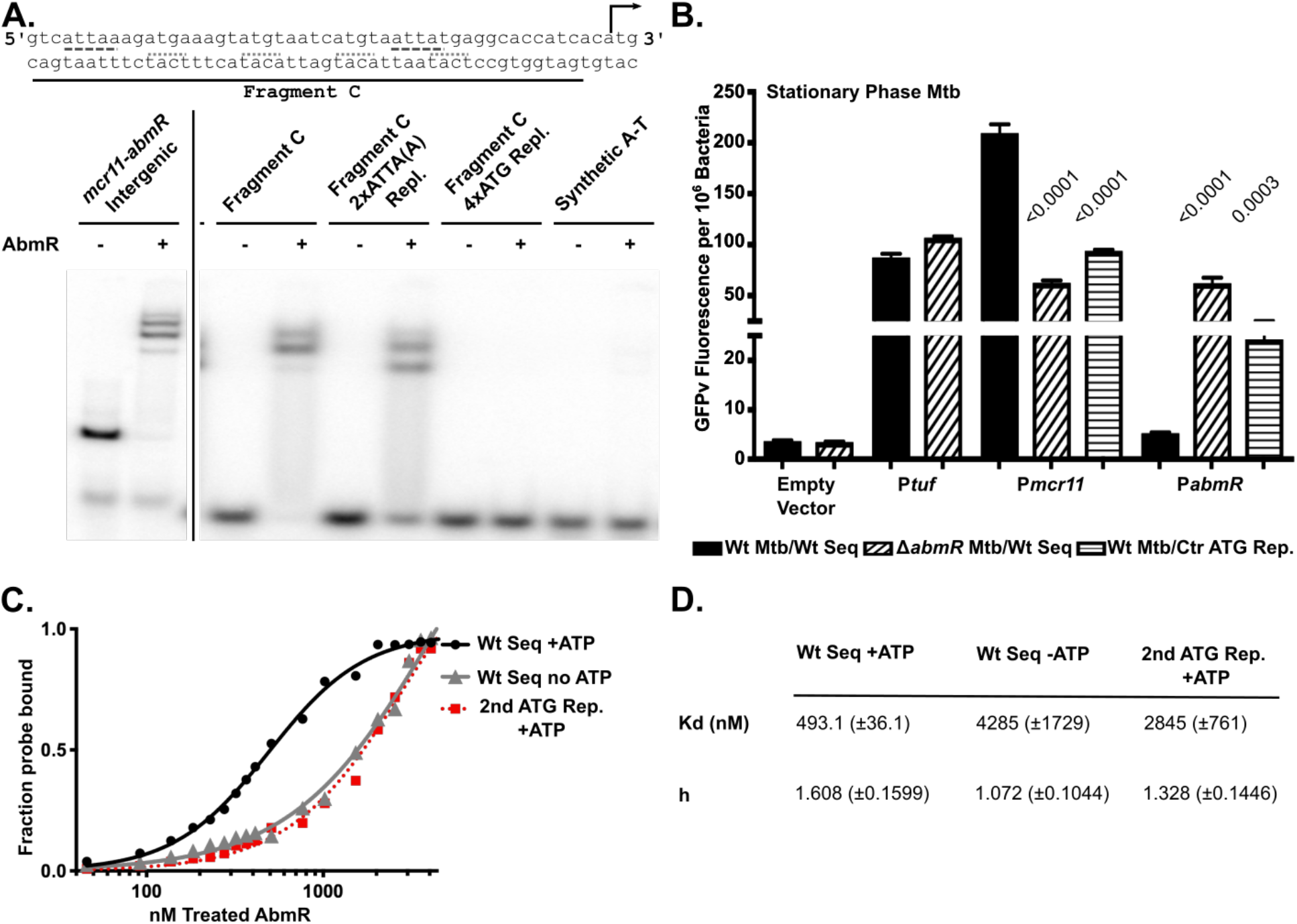
DNA- binding by AbmR is sequence-specific and required for gene regulation. ATP and a unique repeating sequence element contribute to the cooperative binding of AbmR at the *mcr11-abmR* locus. A. Various fragments of DNA were tested for AbmR binding in the presence of ATP by electromobility shift assay (EMSA). Cropped sections of the gel are shown and separated by a black line. B. Stationary phase strains of *Mycobacterium tuberculosis* H37Rv grown in shaking, hypoxic (1.3% O2, 5% CO2) conditions were assayed for promoter;GFPv reporter fusion expression by fluorescence. Expression was normalized to the number of bacteria in each sample. A negative promoterless control and a positive control (elongation factor Tau, *tuf*) were included as assay controls. All statistical comparisons were made between the wild type strain with wild type sequence controls and the test strain using one-way ANOVA with Sidak’s correction for multiple comparisons in Graphpad Prism version 8 software. C. Binding plot of the EMSA results in Figure 1C compared to results in Supplemental Figures 1F and 1G, in which the binding reactions lacked ATP or had a mutated AbmR binding site. D. A table with the details of the Kd and Hill slope for RNase treated AbmR with wild type sequence with or without, and with ATP and the AbmR binding site mutated at the second ATG repeat using a one-site specific binding with Hill slope non-linear regression model in Graphpad Prism version 8 software.

The center two of the four ATGN_5_ repeats were replaced in promoter:GFPv transcriptional fusion reporters of *mcr11* and *abmR* and tested for their impact on the *in vivo* gene-regulatory function of AbmR. In both log- and stationary-phase Mtb, AbmR strongly activated *mcr11* expression and repressed *abmR* expression (Supplemental Figure 1F, Figure 2B). Either deletion of the *abmR* locus or replacement of the center two ATGN_5_ repeats within AbmR’s binding site was sufficient to cause significant loss of *mcr11* promoter activation and de-repression of expression from the *abmR* promoter (Figure 2B). The apparent Kd of de-complexed AbmR for a DNA fragment with the second of the 4x ATGN_5_ repeats replaced was significantly greater than that for the intact sequence (Figure 2C-D, Supplemental Figure 1F). The increase in apparent Kd with the mutant DNA sequence is similar to the increase in apparent Kd for the intact sequence in the absence of ATP, suggesting that both repeating sequence elements and the presence of ATP drive the specific affinity of de-complexed AbmR for a DNA ligand (Supplemental Figure 1G, Figure 2C-D). The requirement for the specific repeating sequence element observed within Mtb, and Hill coefficient greater than 1 supports a role for cooperative binding at the *abmR-mcr11* locus in the gene regulatory function of AbmR.

### RNA-binding by AbmR is promiscuous

A large and diverse array of AbmR co-purifying RNAs demonstrated that AbmR binding of RNA in *E. coli* is promiscuous, in contrast to AbmR’s specific DNA-binding activity. We generated RNA-seq libraries from RNA recovered from purified 39S AbmR complexes and compared them to rRNA-depleted *E. coli* transcription libraries prepared from cells expressing AbmR (Supplemental Table 2). Approximately one-third of transcripts from the control libraries mapped to the AbmR expression plasmid, and the remainder to the genome (Supplemental Tables 2 and 4, Figure 3A). In contrast, nearly 60% of AbmR co-purified transcripts mapped to the AbmR expression plasmid, with the AbmR open reading frame (ORF) being the most abundantly represented transcript (Supplemental Table 4, Figure 3A). Of the 2,590 AbmR co-purified transcripts that mapped to the *E. coli* genome, levels of 985 were significantly different compared to those of the *E. coli* transcriptome control, and these included all categories of transcripts (predicted RNAs, stable RNAs, ribosomal RNAs, tRNAs, and protein coding genes) (Figure 3C, upper panel). 99 of the 985 AbmR co-purified transcripts were enriched >5-fold relative to the control, and these included 30 of the most abundant transcripts that mapped to the *E. coli* genome (Supplemental Table 3, Figure 3B). The 50 most abundant AbmR co-purified RNA samples included only predicted RNAs (56%) and protein-coding genes (44%) (Figure 3C, bottom panel). No specific sequence motifs were identified among the recovered transcripts, so we further characterized the AbmR’s RNA binding properties using a subset of the recovered transcripts, as described below.

**Figure 3:**
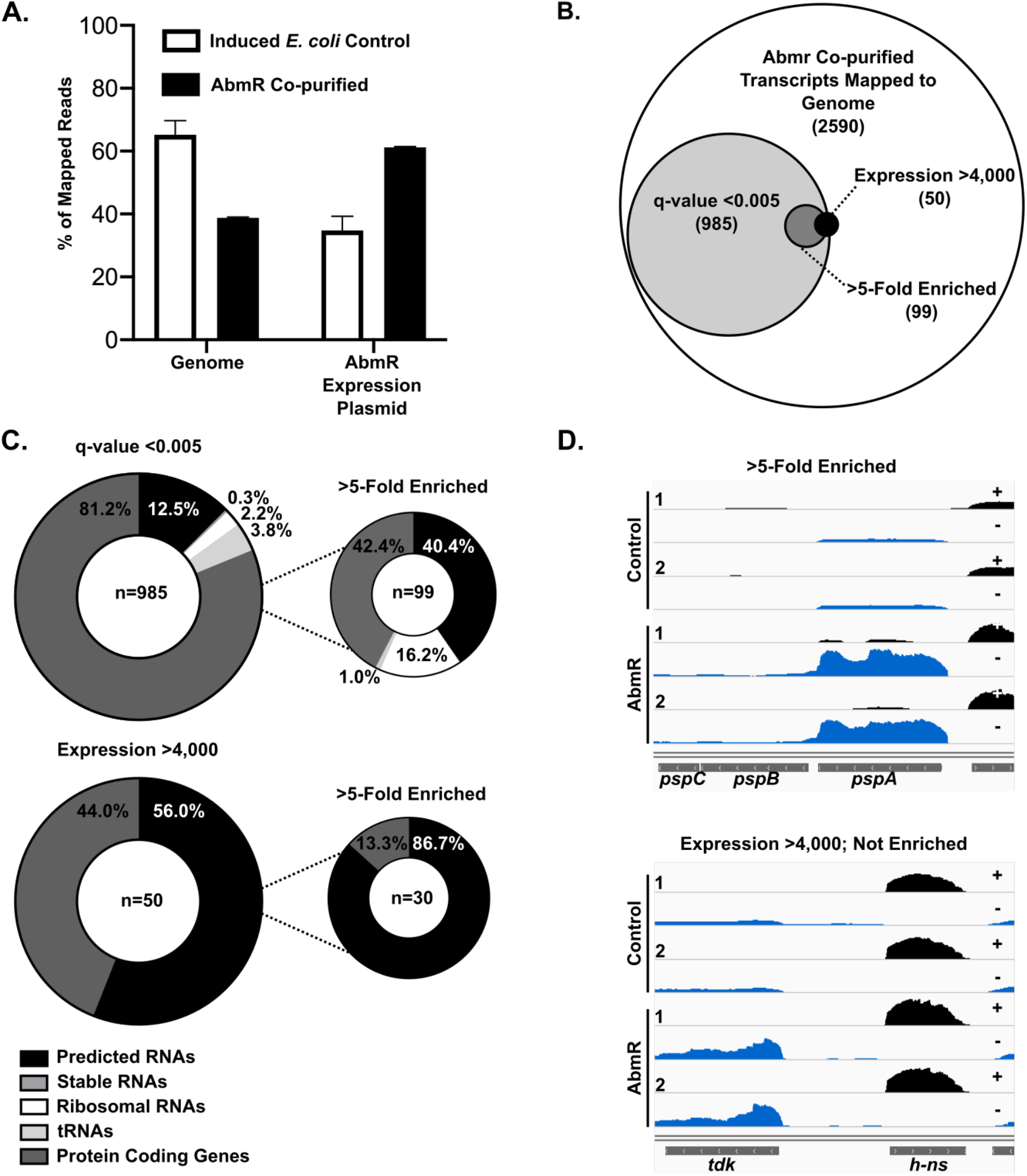
RNA- binding by AbmR is promiscuous. AbmR binding of RNA is promiscuous. A. The fraction of reads that mapped to the genome or the AbmR expression plasmid from induced *Escherichia coli* control RNA-seq reads or from AbmR co-purified RNA-seq reads. All samples had >5 million mapped reads. B. The distribution of AbmR co-purified transcripts that mapped to the genome. C. The distribution of the AbmR co-purified transcripts that mapped to the genome and were significantly different in abundance compared to the induced *E. coli* control (top) or were among the most abundant transcripts detected (bottom). D. Traces of mapped RNA-seq data from both repeats of the induced *E. coli* control (Control) or AbmR co-purified (AbmR) samples. A representative trace from a gene that was >5-fold enriched in the AbmR co-purified samples and a representative trace from a gene that was among the most abundant but not >5-fold enriched are shown.

### AbmR has distinct DNA and RNA-binding determinants

We previously established arginine 146 (R146) as a critical determinant of AbmR’s DNA-binding activity (10), and reconfirmed its importance for DNA binding by using de-complexed R146A AbmR protein (Figure 4A). However, AbmR R146A formed RNAase A-sensitive, DNase I-resistant 39S complexes that were indistinguishable from those of Wt AbmR and it co-purified with RNA (Supplemental Figure 2A). These results show that the R146A mutation differentiates between RNA versus DNA binding (Figure 4B), so we tested the ability of Wt versus the R146A mutant AbmR protein to bind individual RNA transcripts identified in the AbmR co-purification experiment.

**Figure 4:**
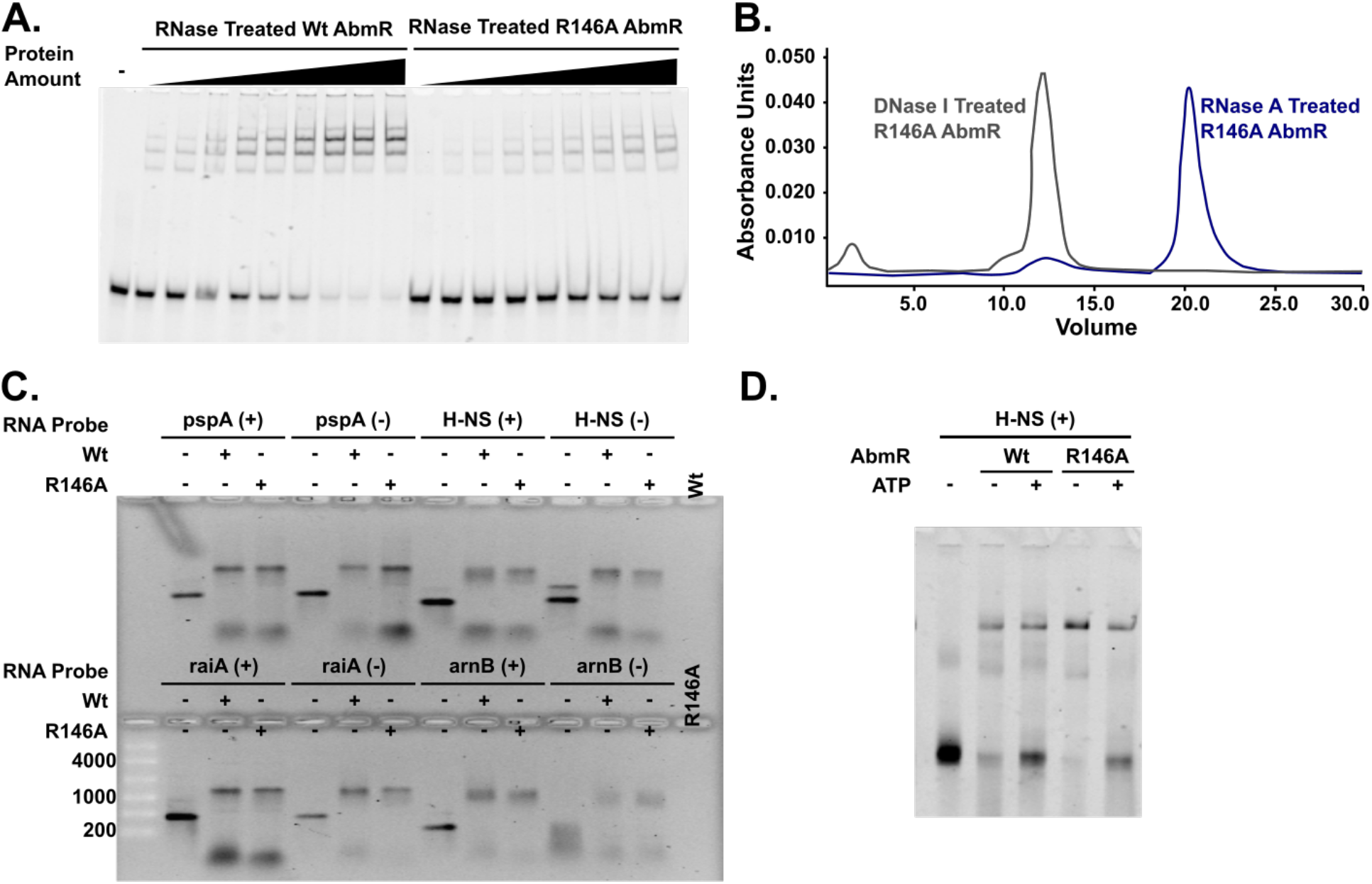
AbmR has distinct DNA- and RNA-binding determinants. AbmR RNA-binding is resistant to mutation of R146A in the putative nucleic acid binding domain of AbmR and ATP independent. A. The DNA binding activities of de-complexed Wt and R146 AbmR was tested with the *mcr11/abmR* promoter DNA fragment with ATP by electromobility shift assay (EMSA). B. Size exclusion chromatograms of R146A AbmR treated with DNaseI or RNase A reveal similar size profiles to wild type (Wt) AbmR. C. *In vitro* transcribed RNA from the coding (+) or non-coding (-) strands of several RNA-seq hits was mixed with equal amounts of de-complexed Wt or R146A AbmR in the presence of ATP and loaded into an ethidium bromide stained EMSA. The RiboReady 1 kb denatured single stranded RNA ladder is shown for size approximation. Protein only controls were loaded to demonstrate that de-complexed AbmR had no residual UV trace. D. Fluorescein-UTP tagged *in vitro* transcribed RNA from the coding (+) strand of *h-ns* was added to limiting but equivalent amounts of de-complexed Wt or R146A AbmR in the absence or presence of ATP and resolved by EMSA.

The *abmR* ORF and six additional protein coding transcripts were selected from the RNA-seq experiment for analysis, including: transcripts that differed significantly between the *E. coli* transcriptome control and AbmR co-purified RNA samples; highly abundant transcripts in the AbmR co-purified RNA samples; and transcripts that were >5-fold enriched in the AbmR co-purified RNA samples (Supplemental Table 5, Figure 3D). The sRNA Mcr11 was also chosen for its relevance to the established *in vivo* regulatory functions of AbmR.

De-complexed wild type (Wt) and R146A AbmR proteins were tested for their ability to bind the *in vitro* transcribed RNAs. We found that Wt and mutant AbmR were similarly competent for RNA-binding in the presence of ATP (Figure 4C, Supplemental Figure 2B). In addition, RNA-binding by de-complexed AbmR was comparable across all ligands identified from the RNA-seq experiment, regardless of whether they came from the >5-fold enriched group or because they were among the most abundant transcripts in the AbmR co-purified libraries.

The sequence specificity of AbmR’s RNA-binding function was tested by comparing the *in vitro* RNA-binding activity of de-complexed AbmR for RNA ligands transcribed from both the sense (+) and anti-sense (-) strands of the six *E. coli* protein-coding RNA ligands selected for validation. Binding of sense and antisense *in vitro*-transcribed RNA ligands was similar for both de-complexed Wt and the R146A mutant AbmR proteins (Supplemental Figure 2B, Figure 4C). Additionally, de-complexed Wt and the R146A mutant AbmR proteins were competent for RNA-binding in the presence or absence of ATP (Figure 4D). While de-complexed AbmR also robustly bound an *in vitro*-transcribed AbmR ORF RNA ligand with little to no dependence on ATP, binding of *in vitro-*transcribed Mcr11 was only marginal (Supplemental Figure 3A). Thus, while de-complexed AbmR can bind RNA promiscuously, there may be a bias toward some RNA ligands, for reasons yet to be determined. Together, these results indicate that de-complexed AbmR is capable of non-specifically binding RNA ligands from *E. coli* and Mtb in an R146-independent manner and they establish that the DNA- and RNA-binding determinants of AbmR are distinct from one another.

### AbmR has an RNA-protective function

Having established that AbmR is capable of binding both RNA and DNA, we next determined the preferred ligand of AbmR. Competitive binding reactions with labelled, *in vitro*-transcribed *h-ns* RNA were set up in the presence of ATP and 50-1,000 fold excess unlabelled RNA or dsDNA (containing AbmR binding sites. De-complexed Wt or R146A AbmR was added to the binding reactions with both ligands present and the competition reactions were resolved by EMSA. In the ligand competition assay, de-complexed AbmR preferentially bound the DNA ligand in an R146A-dependent fashion (Figure 5A). Excess unlabelled RNA was not able to displace bound, labelled RNA ligand(s), suggesting that once bound, the RNA binding domain of AbmR and the bound RNA ligand may be relatively inaccessible to competing RNAs. This result is concordant with our previous finding that AbmR complexes purified from *E. coli* did not have detectable RNA-binding activity when provided with *in vitro*-transcribed RNA (10). In the absence of an added RNase inhibitor, residual RNase A in the de-complexed AbmR protein preparations degraded unbound RNAs (Figure 4D). While the presence of ATP had the expected effect of decreasing the activity of the residual RNase A in the binding reactions (Figure 4D) (21), these observations prompted us to investigate the ability of AbmR to protect RNA ligands from limited RNase A-digestion in the absence of ATP.

**Figure 5:**
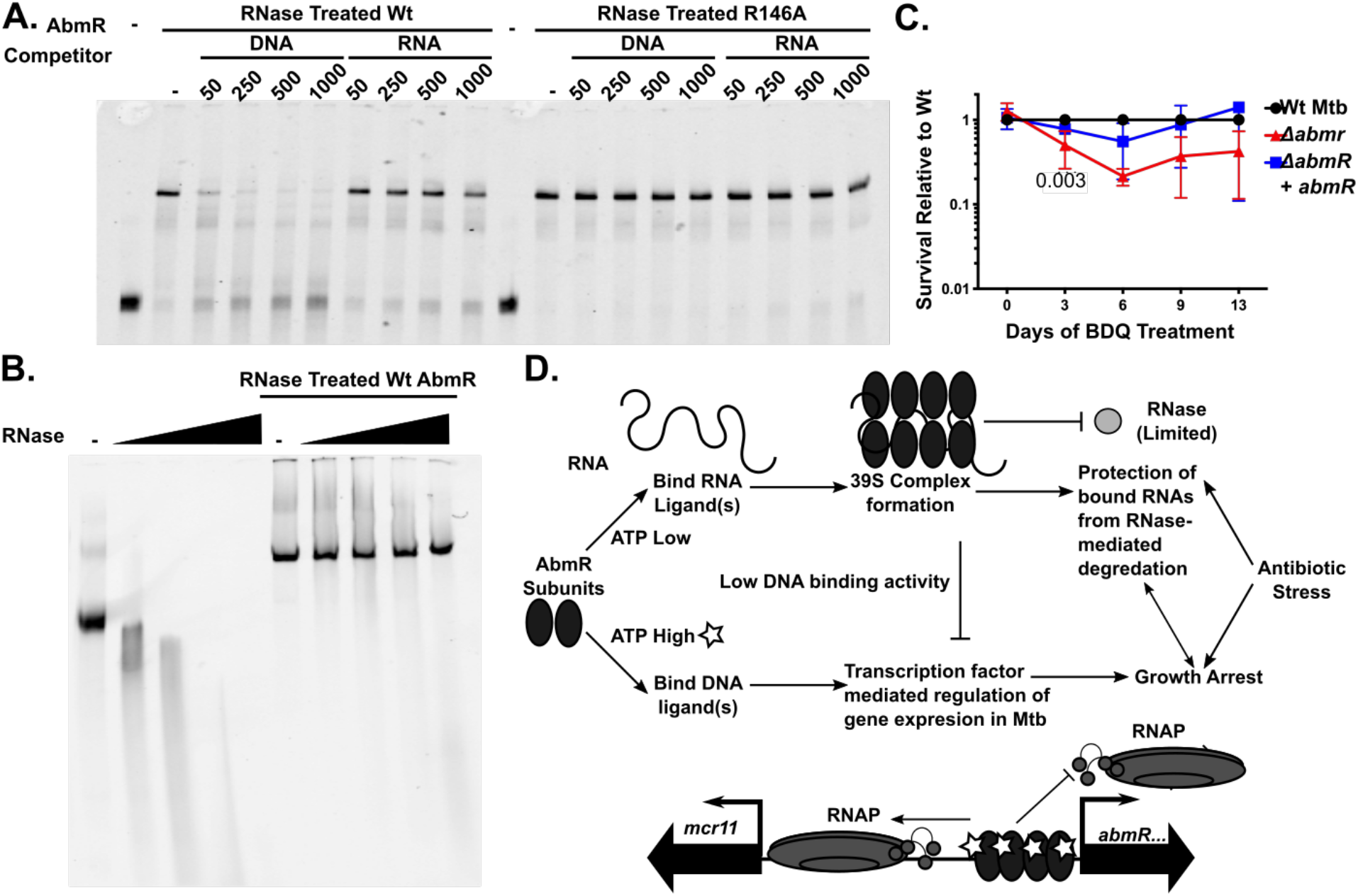
AbmR has RNA protective activity. AbmR binds DNA preferentially and has an RNA protective function. A. Equivalent amounts of de-complexed wild type (Wt) or R146A were added to a mixture containing fluorescein-UTP tagged *in vitro* transcribed RNA from the coding (+) strand of *h-ns* and the indicated fold-excess of unlabelled competitors in the presence of ATP and absence of murine RNase inhibitor, then resolved by electromobility shift assay (EMSA). B. Fluorescein-UTP labelled, *in vitro* transcribed AbmR ORF RNA was added to increasing amounts of RNaseA in the presence or absence of de-complexed Wt AbmR with a limited quantity of murine RNase inhibitor and visualized in an EMSA. C. *Mycobacterium tuberculosis* H37Rv Wt, *ΔabmR* or *ΔabmR* complemented strains were treated with 20X the minimum inhibitory concentration of bedaquiline or DMSO vehicle control and bacterial survival was assayed by plating CFUs. Relative survival was calculated by dividing CFUs from treatment time points by starting CFU inputs and ratioing to the Wt strain. The results are the means of three independent repeats. A two-way ANOVA with Bonferroni correction for multiple comparisons was used to assess significance. D. A schematic model showing the various roles of AbmR’s DNA and RNA-binding functions within the mycobacterial cell. AbmR protein subunits are shown as filled ovals, ATP molecules as open stars, genes on the chromosome as dark, filled arrows, and transcriptional start sites are represented as thin arrows. Cartoons shown within the schematic represent RNA polymerase (RNAP), RNA, and RNase, as labelled.

The minimal quantity of murine RNase inhibitor required to block residual RNase activity in de-complexed AbmR protein preparations was determined (data not shown). An RNase protection assay was then performed using this quantity of murine RNase inhibitor, an RNA ligand, de-complexed Wt or R146A AbmR proteins, and increasing amounts of RNase A. De-complexed Wt or R146A AbmR were able to protect bound RNA ligands from levels of RNAse A that completely degraded unbound RNA in the absence of ATP (Figure 5B, Supplemental Figure 3B). RNA ligands that were protected from degradation included extracted, co-purifying RNA from *E. coli* as well as the AbmR ORF mRNA from Mtb. Together, these data show that AbmR can protect bound RNAs from limited RNase digestion in the absence of ATP.

### AbmR is required for optimal survival of bedaquiline treatment

The antimycobacterial drug bedaquiline targets the activity of the ATP synthase machinery and causes a dramatic depletion in cellular ATP pools, delayed bacterial death, and enhanced RNA stability (22,23). The expression of *abmR* has been reported to significantly increase in Mtb exposed to bedaquiline or the frontline drug isoniazid (15). Thus, we tested the expression of the *mcr11* and *abmR* loci in response to treatment with bedaquiline, isoniazid, or a DMSO vehicle control using promoter:GFPv transcriptional fusion reporters in Mtb. Expression of *mcr11* was unaltered by exposure to isoniazid, but significantly up-regulated in response to bedaquiline treatment at early and later time points (Supplemental Figure 3C). Promoter activity at the *abmR* locus trended upward in response to both isoniazid and bedaquiline treatment, but this trend did not reach statistical significance (Supplemental Figure 3C). Survival of Wt, *ΔabmR*, and *ΔabmR* complemented strains of Mtb after exposure to bedaquiline was monitored by CFU plating over two weeks. Three days post-treatment, survival of the *ΔabmR* strain was significantly lower than Wt or complemented Mtb, and the trend of reduced survival of this strain was observed for the remainder of the study period (Figure 5C). These results indicate a positive role for *abmR* in the optimal survival of Mtb exposed to bedaquiline.

## DISCUSSION

These results establish AbmR as a dual function transcription factor that alternates between its low molecular weight DNA binding form and large 39S ribonucleoprotein complexes. We found that AbmR has distinct DNA- and RNA-binding determinants and that RNA binding stabilizes the 39S AbmR complexes. RNA binding was promiscuous, yet AbmR-associated RNA was protected from degradation. The biological importance of RNA binding for AbmR remains to be determined, but our findings are consistent with a role in Mtb’s adaptation to stress.

### AbmR binds RNA promiscuously and has an RNA-protective function

AbmR is critical for the growth-phase dependent activation of *mcr11* expression at the levels of both transcription and stabilization, although the stabilizing mechanism remains unclear (10,11,16). In *E. coli* and a wide variety of pathogenic bacteria, the hexameric RNA-binding protein Hfq facilitates the antisense interaction between sRNAs and their regulatory targets (24). ProQ/FinO are a more recently appreciated family of RNA chaperones that also bind the 3’ end of sRNAs and modulate their interactions with mRNA targets (25,26). While Mtb lacks both Hfq and ProQ/FinO orthologs (27), the weak interactions between AbmR and *in vitro*-transcribed Mcr11 fail to support a model in which AbmR serves as an Mcr11 or global sRNA chaperone. Rather, AbmR appears to more generally sequester and protect bound RNAs from degradation.

Global stabilization of transcripts is a generalized stress response that occurs in Mtb and the model organism Msm (18,23,28). The mechanisms that regulate RNA stability are complex and intersectional. They include transcription and/or translation rates of the RNA, the presence of RNA secondary structures, post-transcriptional modifications of the RNA, binding of RNA chaperones or other proteins, and the abundance or access of RNA-degrading enzymes to the transcript. A recent evaluation of mycobacterial RNA degradasome components in nutrient-replete bacteria revealed expected components as well as cold-shock proteins and novel KH-domain proteins (29). The impact of stress on specific mycobacterial degradasome components and other regulators of RNA stability is not well characterized. Recent work has demonstrated that RNA stability in Mtb is responsive to antibiotic treatment that alters the metabolic status of the cell and not simply explained by the abundance of the transcript or RNase levels (23). The AbmR requirement for optimal survival of bedaquiline treatment suggests the protein has a role in stress response and adaptation of Mtb to this ATP-depleting drug. While the abundance of co-purifying transcripts from the AbmR over-expression plasmid suggests that abundance and proximity are key determinants of the RNA ligand selection by AbmR, the relative importance of these and other physiological factors is not yet clear. One limitation of the present study is that AbmR’s RNA binding ligands were profiled in an *E. coli* expression background. Efforts are underway to determine whether AbmR has more preferred and/or biologically relevant RNA targets within a native Mtb environment. Further investigation of the role(s) of AbmR’s regulatory functions and impacts on RNA stability will be important for understanding stress tolerance in Mtb, including that caused by antibiotic treatment.

### DNA-binding by AbmR is cooperative and sequence specific

In contrast to its RNA interactions, AbmR binding to DNA is sequence specific, and shows cooperativity for binding at the operator sequences that regulate expression of *mcr11* and *abmR*. AbmR recognizes an octameric direct repeat sequence of ATGN5 that occurs four times in tandem at the *mcr11-abmR* locus, and EMSAs show 4 distinctly migrating complexes. Our working hypothesis is that these discrete complexes are formed by AbmR progressively binding to each of the four tandem ATGN_5_ sites within the DNA, such that binding to the second, third and fourth sites is facilitated by AbmR-AbmR interactions at adjacent sites.

Our finding that treatment of the 39S complexes of AbmR with RNase increases the DNA-binding affinity by 100-fold supports a model in which the binding sites for RNA and DNA are overlapping and DNA is preferentially bound by the decomplexed form of AbmR. Despite overlap, the binding determinants for DNA and RNA are not identical, as Arginine 146 is not required for RNA binding or 39S complex formation, but strongly enhances DNA binding. Moreover, ATP enhances AbmR binding to DNA ∼10-fold but does not stimulate binding to RNA. This differential effect of ATP on nucleic acid binding distinguishes between AbmR’s binding with DNA versus RNA, and the physiological significance of ATP for AbmR’s regulatory activities warrants further investigation. While ATP does not appear to affect RNA binding directly, the level of ATP could regulate the predominant activity of AbmR in cells: when ATP levels are high, AbmR would be predicted to preferentially bind DNA; when ATP levels are low, AbmR would have greater opportunity to bind RNA and form 39S complexes.

AbmR is distinct from previously characterized dual-function nucleic acid binding proteins in bacteria such as DNA-binding protein from starved cells (Dps) proteins and nucleosome associated proteins (NAPs). Dps proteins are expressed during stationary phase and form high order oligomers that present as large, distorted icosahedrons that have storage and antioxidant functions (30). NAPs organize and compact the genome, while also influencing gene expression through transcription factor activity (31). The 39S complexes formed by AbmR do not resemble the higher order DNA binding structures formed by NAPs in that they preferentially bind RNA and they lack evidence of the ferroxidation centers associated with Dps proteins (32,33). However, several NAPs bind both DNA and RNA. Histone-like nucleoid structuring (H-NS) protein, which is found in *E. coli* and related bacteria binds AT rich sequences on DNA and forms gene-silencing filaments through head-to-head and tail-to-tail interactions between neighbouring H-NS subunits (34). Additionally, H-NS is capable of specifically binding both mRNA and sRNA and decreasing the *in vitro* and *in vivo* stability of the bound transcripts (35). Lsr2 is a functional ortholog of H-NS in Mtb that forms compacted nucleoids after binding AT-rich regions of Mtb’s genome (36). Lsr2 is essential for hypoxic adaptation and virulence, but it has not been reported to bind RNA or alter RNA degradation (37,38). AbmR’s DNA-binding motif is AT-rich like those bound by some NAPs, but it is also a specific tandem repeat that occurs only once in Mtb’s genome. Elucidation of the architecture of how AbmR is arranged along its bound operator DNA may yield information on whether local compaction of DNA contributes to AbmR’s biological roles in Mtb. AbmR also has some functional similarity to the SarA family of winged helix motif-containing transcription factors that regulate virulence in *Staphylococcus aureus*. One of these, SarA, regulates gene expression as a transcription factor, but also binds and stabilizes a specific subset of virulence-associated RNAs *in vivo* (39). In contrast, AbmR’s RNA-binding activity appears to be relatively non-specific and SarA is not known to form AbmR-like complexes.

### Working Model

We present a working model based on these findings, in which distinct DNA-binding and RNA-binding determinants allow AbmR to regulate gene expression, modulate RNA stability, and facilitate bacterial adaptation to stress in unexpected ways. We propose that in ATP-rich environments, non-complexed AbmR preferentially binds a unique sequence element at the *mcr11-abmR* locus, activating *mcr11* expression and repressing *abmR* expression like a classical bacterial transcription factor (Figure 5D, bottom). Mcr11 could then accumulate in a growth phase-dependent manner, contributing to growth arrest in Mtb. In low/depleted ATP conditions, decreased binding of AbmR to DNA would de-repress *abmR* expression. Abundant, non-complexed AbmR could then bind proximal RNA ligands and sequester them from degradation in the 39S complex, feeding back into de-repression of *abmR* (Figure 5D, top). This is a complex auto-regulatory system mediated by competition between the DNA and riboregulatory RNA-bound states of AbmR. While physiological targets of AbmR’s RNA regulatory activity have not been defined, the RNA-stabilized phenotype is associated with conditions that induce growth arrest, non-replicating persistence, and antibiotic tolerance. AbmR may contribute to growth-arrest and stress tolerance through the regulation of the sRNA Mcr11, as well as through modulation of RNA ligands sequestered in the 39S complex.

Many unique aspects of AbmR dual-functions remain to be elucidated, including its ability to use distinct binding modalities for DNA and RNA-ligands and forming an RNA regulatory 39S complex. While AbmR orthologs are widely distributed across the genomes of Actinobacteria, information about their conserved functions and biological significance is lacking. A detailed analysis of the structure of AbmR in complex with one or more ligands will facilitate understanding of how AbmR transitions between its mutually exclusive DNA- and RNA-bound states. Similarly, exploration of the biological roles of AbmR’s newly established RNA-binding function will advance understanding of the stress tolerance pathways of one of the world’s most deadly pathogens.

## AVAILABILITY

Inkscape is freely available (https://inkscape.org/)

Rockhopper is freely available (https://cs.wellesley.edu/~btjaden/Rockhopper/)

RNA-seq data has been deposited to the Sequence Read Archive at the National Library of Medicine under BioProject PRJNA738636: AbmR Co-Purifying RNAs from induced *E. coli*.

## SUPPLEMENTARY DATA

Supplementary Data are available at NAR online.

## FUNDING

This work was supported in part by the National Institutes of Health [grant numbers R01AI063499 to KAM; R01GM080573 to JP; and the Potts Memorial Foundation (RG). Funding for open access charge:.

## ACKNOWLEDGEMENT

We thank the Tissue Culture and Media Core at Wadsworth Center for the provision of reagents, and the Applied Genomics Technologies Core for assistance with DNA sequencing, the Biochemistry and Immunology core for assistance with size exclusion chromatography, and the Electron Microscopy core for imaging. We are very grateful to Joseph Wade and Erica Lasek-Nesselquist for technical assistance and guidance.

## CONFLICT OF INTEREST

The authors declare that they have no conflicts of interest.

## Notes

### Competing Interest Statement

The authors have declared no competing interest.

