## Supplemental figures for "AbmR is a mycobacterial dual-function transcription factor and ribonucleoprotein with distinct DNA and RNA-binding determinants"

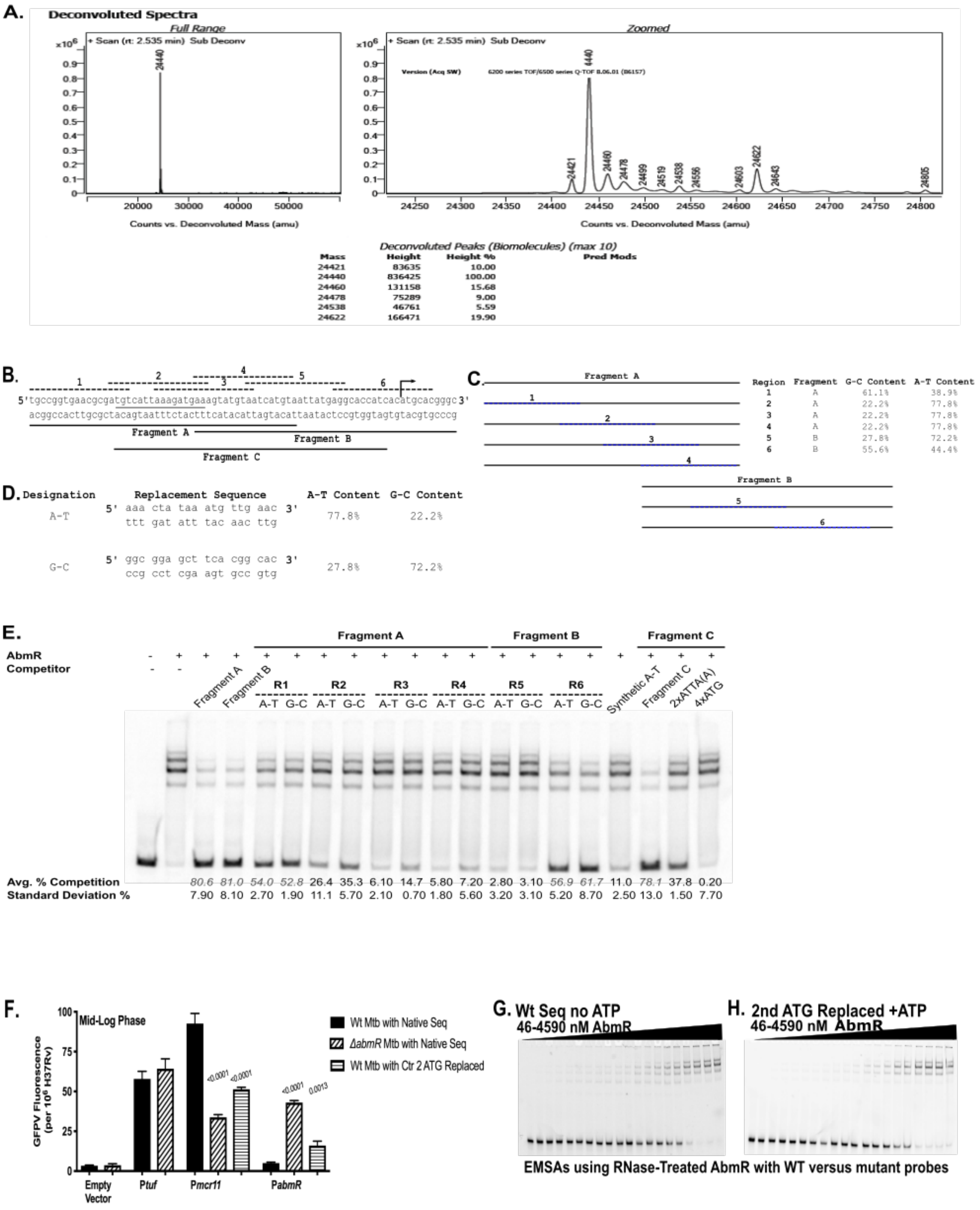

A.
Deconvoluted Spectra
Full Range
Zoomed
+ Scan (rt: 2.535 min) Sub Deconv
Version (Acq SW) 6200 series TOF/6500 series Q-TOF B.06.01 (B6157)
Counts vs. Deconvoluted Mass (amu)
Deconvoluted Peaks (Biomolecules) (max 10)
Mass Height Height % Pred Mods
24421 83635 10.00
24440 836425 100.00
24460 131158 15.68
24478 75289 9.00
24538 46761 5.59
24622 166471 19.90
B.
5'tgccggtgaacgcgatgtcattaaagatgaaagtatgtaatcatgtaattatgaggcaccatcacatgcacgggc3'
acggccacttgcgctacagtaatttctactttcatacattagtacattaatactccgtggtagtgtacgtgcccg
Fragment A
Fragment B
Fragment C
C.
Region Fragment G-C Content A-T Content
1 A 61.1% 38.9%
2 A 22.2% 77.8%
3 A 22.2% 77.8%
4 A 22.2% 77.8%
5 B 27.8% 72.2%
6 B 55.6% 44.4%
D.
Designation Replacement Sequence A-T Content G-C Content
A-T 5' aaa cta taa atg ttg aac 3' ttt gat att tac aac ttg 77.8% 22.2%
G-C 5' ggc gga gct tca cgg cac 3' ccg cct cga agt gcc gtg 27.8% 72.2%
E.
AbmR
Competitor
Fragment A
Fragment B
Fragment C
R1 R2 R3 R4 R5 R6
A-T G-C
Synthetic A-T
2xATTA(A)
4xATG
Avg. % Competition 80.6 81.0 54.0 52.8 26.4 35.3 6.10 14.7 5.80 7.20 2.80 3.10 56.9 61.7 11.0 78.1 37.8 0.20
Standard Deviation % 7.90 8.10 2.70 1.90 11.1 5.70 2.10 0.70 1.80 5.60 3.20 3.10 5.20 8.70 2.50 13.0 1.50 7.70
F.
Mid-Log Phase
GFPV Fluorescence (per 10⁶ H37Rv)
Wt Mtb with Native Seq
ΔabmR Mtb with Native Seq
Wt Mtb with Ctr 2 ATG Replaced
Empty Vector
Ptuf
Pmcr11
PabmR
<0.0001
0.0013
G. Wt Seq no ATP
46-4590 nM AbmR
H. 2nd ATG Replaced +ATP
46-4590 nM AbmR
EMSAs using RNase-Treated AbmR with WT versus mutant probes

**Supplemental Figure 1**. RNase treated AbmR presents as an intact mass of expected molecular weight and binds DNA in a sequence specific manner. A. Mass-spec showing that RNase-treated and re-purified AbmR presents as an intact mass of expected molecular weight. B. The *mcr11-abmR* locus with DNA sequence elements defined. C. Details of the scanning replacement probes used for competition experiments. D. Details of the AT and GC rich scanning replacement sequences used for competition experiments. E. Scanning competition EMSA narrows the AbmR binding region to a specific area and a repeating sequence element, independent of the AT content of the DNA fragment. F. Promoter activity in log phase Mtb grown in shaking, hypoxic conditions. N = 3 independent biological replicates. Statistical significance assessed with one-way ANOVA using Sidak's correction for multiple comparisons. All comparisons are to the Wt Mtb with Wt sequence reference strain. G. EMSA with RNase A treated and re-purified AbmR protein, the wild type sequence at the *mcr11-abmR* locus, and no added ATP. H. EMSA with RNase A treated and re-purified AbmR protein, the 2nd ATG replaced in the repeating AbmR binding sequence at the *mcr11-abmR* locus, and with 1mM ATP.

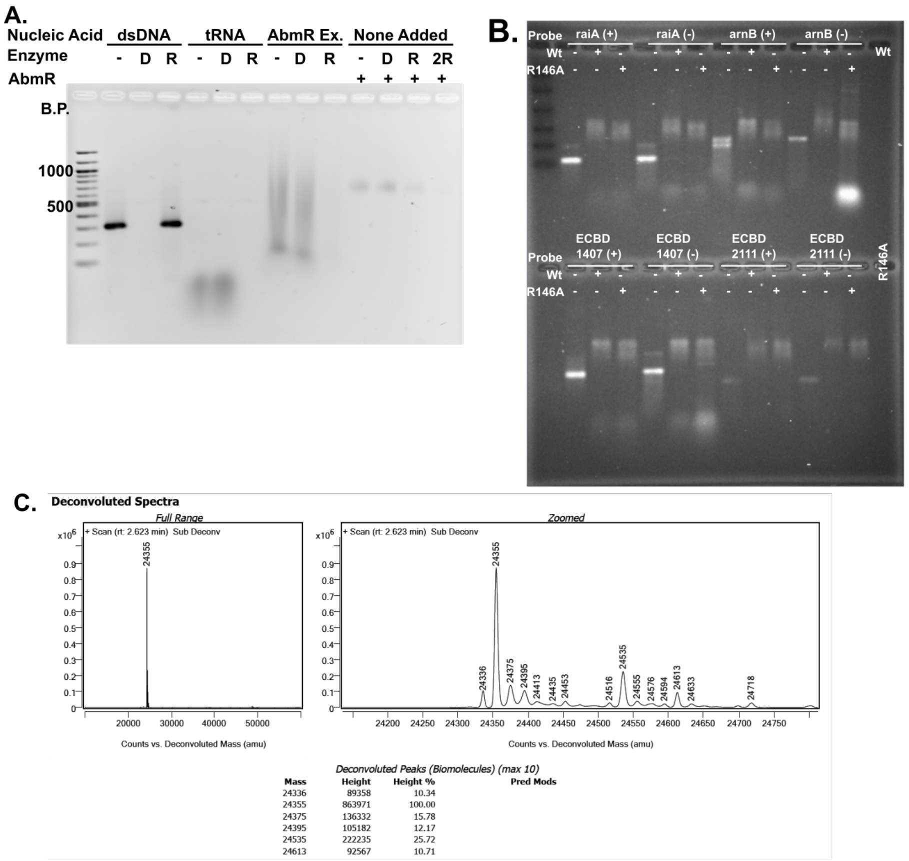


**Supplemental Figure 2.** R146A AbmR has a similar RNA-binding profile to Wt AbmR. A. Ethidium bromide stained agarose gel of the DNase (D) or RNase (R) digestion of double-stranded DNA (dsDNA), tRNA, phenol-chloroform nucleic acid extracted from purified R146A AbmR 39s complexes, (AbmR Ex.) or R146A AbmR 39s complexes with no added nucleic acid (None Added). A 100 base pair dsDNA ladder is loaded for comparison. B. RNA-binding EMSA of Wt AbmR vs R146A AbmR of 8 different ligands show that the proteins have similar RNA-binding function in the presence of ATP and the absence of murine RNase inhibitor. C. Mass-spec of RNase A-treated and re-purified R146A AbmR shows an intact mass of the expected size.

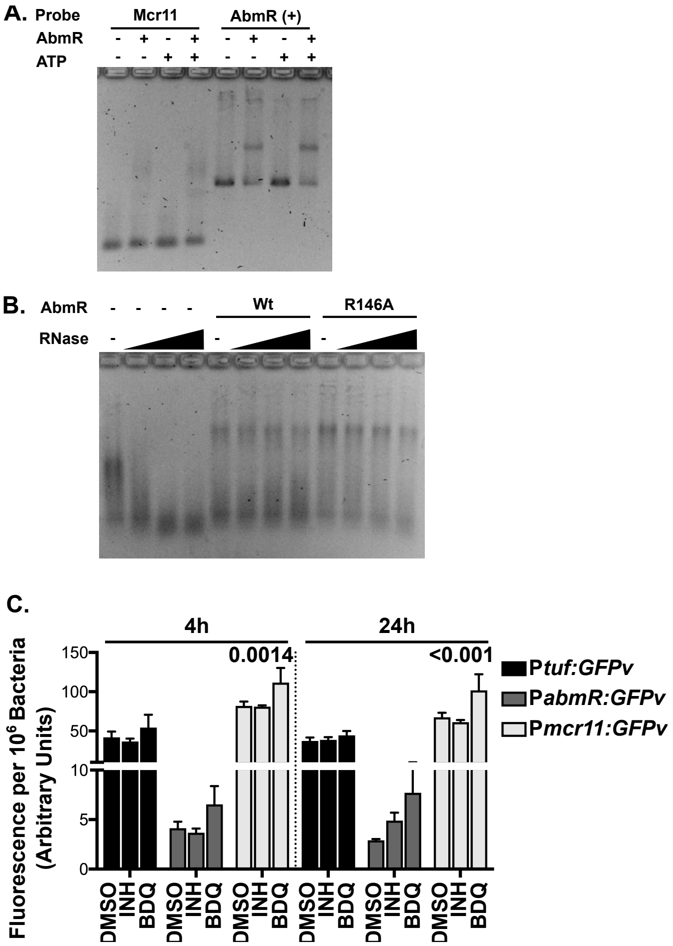


**Supplemental Figure 3.** A. Ethidium bromide stained electromobility shift assay (EMSA) of Wt RNase-treated and re-purified AbmR with *in vitro* transcribed Mcr11 or the AbmR open reading frame without or with 1 mM ATP and murine RNase inhibitor. B. RNA that co-purified with Wt AbmR 39s complexes was extracted and purified. Extracted, co-purifying RNA was added to increasing amounts of RNaseA in the presence or absence of equivalent amounts of RNase-treated and re-purified Wt and R146A AbmR and a limited quantity of murine RNase inhibitor then loaded into an ethidium bromide stained EMSA. C. Expression of promoter:GFPv reporter fusions in Mtb treated with vehicle control (DMSO), isoniazid (INH), or bedaquiline (BDQ). A two-way ANOVA with Bonferroni correction for multiple comparisons was used to assess significance.
